# Douglas fir and Norway spruce admixtures to beech forests along in Northern Germany – Are soil nutrient conditions affected?

**DOI:** 10.1101/2020.09.25.313213

**Authors:** Estela Covre Foltran, Christian Ammer, Norbert Lamersdorf

## Abstract

The establishment of mixed forest stands can be seen as an option to improve soil nutrient conditions and to protect forest ecosystems from various impacts of climate change. Our study analyzed groups of pure mature European beech (*Fagus sylvatica*; B), Douglas fir (*Pseudotsuga menziesii*; D) and Norway spruce (*Picea abies*; S) stands as well as mixtures of beech with either Douglas fir (DB) or spruce (SB), i.e. 5 forest stands per site along Northern Germany with two regionally clearly differing sites conditions (i.e., 3 loess-influenced, loamy sites in the Solling region, southern Lower Saxony and 4 sandy lowland sites further north). In order to determine possible influences of the tree species and their mixtures on soil properties, the organic layer and the mineral topsoil were first chemically characterized for all 35 plots down to a depth of 30 cm (pH, C, N, P, CEC, exchangeable nutrient cation contents and stocks, base saturation-BS). Our results indicated, independent of sites condition, that pure S stands showed the lowest pH and BS, meanwhile B the highest BS. The impact of D varied depending on site condition. On sandy soils, pure D showed higher pH and BS than under pure S, while on loamy soils the pH under D and B was lower than under S. Regarding cations stocks under sandy soils conditions, S stands and its admixture SB depleted soil Ca and Mg stocks more than pure D and B. In contrast, under loamy soil conditions B showed depleted (lowest) soil exchangeable Ca and Mg stocks more than under S stands. Soil exchangeable K under mixed stands were among the highest compared to pure stands, independent of the site condition. Thus, mixed species stands generally decreased soil base cation depletion compared to pure conifer stands. Admixtures of Douglas-fir (DB) seem to lead to smaller changes in pH, CEC and BS than those of Norway spruce, this effect become more important at sandy soil sites. Therefore, forest management may consider mixtures of European beech and Douglas fir as a reasonable management option without apprehending negative effects on soil chemistry.

## 1. Introduction

The development of forest soils is a complex process driven by abiotic and biotic factors (Binkley and Giardina, 1998). The existing geologic parent material, climate conditions and the topography of a given site are essential to formation of forest soils (Hansson et al., 2020), however, those key characteristics develop very slowly. Much faster processes caused by presence (or absence) of particular tree species are known to alters the development of soils in major ways, at times scales of decades (Finzi et al., 1998; van Breemen et al., 1997). Biotic factors are critical to forest soil formation, impacting soil biological, soil physical and soil chemical processes and characteristics (Dawud et al., 2017; Vesterdal and Raulund-Rasmussen, 1998). Thus, site specific forest covers potentially leads to distinct impacts on forest soil chemical and soil physical processes, as well as on soil biodiversity.

Compositionally and structurally diverse forests represent an important element of approaches to deliver a wide range of ecosystem goods and services (Cremer and Prietzel, 2017). The establishment and management of mixed stands is discussed as an effective measure to adapt forests stands to climate change (Ammer et al., 2008; Neuner et al., 2015) and other global challenges such as air pollution and invasive species (Bauhus et al., 2009).

In Central Europe enrichments of European beech (*Fagus sylvatica*) stands with native Norway spruce (*Picea abies*) or non-native Douglas fir (*Pseudotsuga menziesii menziesii*), may result in mixtures that provide income and cope better with the envisaged hazards (Neuner et al., 2015). However, reports of frequent droughts, windthrow and bark beetle infestations around Europe, induced by climate change, make the wide-spread use of native conifers, e.g. Norway spruce (*Picea abies*) in Central Europe, increasingly problematic since the is heavily affected by these hazards (Dobor et al., 2020; Hlásny and Turčáni, 2013; Kölling and Zimmermann, 2007). Thus, coastal Douglas fir is considered a suitable alternative forest tree species to Norway spruce. Douglas fir is characterized by fast growth, good wood features and a high tolerance to heat and drought, which makes it a highly profitable tree species at appropriate sites across Europe (Kownatzki et al., 2011).

Ecological characteristics of mixed species stands are often intermediate in comparison with pure stands of the corresponding species (Augusto et al., 2015; Rothe and Binkley, 2001). Nutrient facilitation process through complementary effects of the different tree species in mixed stands have been reported by several authors (Comerford et al., 2006; Foster and Bhatti, 2006; Lambers et al., 2008; Rakshit et al., 2015; Schmidt et al., 2015) and may explain higher productivity when compared with pure stands in some cases (Ammer, 2019). Cremer and Prietzel (2017) investigated mixed forest effects on mineral soil base saturation and pH and concluded that overall tree species mixtures appeared to improve soil base cation stocks. However, mixture effects on forest soil chemistry vary depending on tree species identity, climatic factors and soil type (Augusto et al., 2015).

From a management point of view the selection of tree species with desired characteristics, e.g. complementary traits for resource use, is one of the most important silvicultural decisions (Schall and Ammer, 2013). However, creating mixtures depending on the soil chemical status requires careful tree species selection rather than increasing tree species diversity *per se* (Dawud et al., 2017). Conifers are known to increase C stocks, while many broadleaves species are able to increase base saturation at top-mineral soil (Cremer and Prietzel, 2017). The impact of Douglas fir on biogeochemical cycles have been extensively studied (Marques et al., 1997; van Miegroet and Cole, 1985; Zeller et al., 2019). However, not much is known how mixtures of non-native Douglas fir and native European beech interact and shape soil chemistry. Due to the high fine root density in deeper soil layers observed under Douglas fir (Calvaruso et al., 2011), decreases on nutrient leaching and cation losses might be expected and would differ from pattern that have been observed under the native conifer Norway spruce (Oulehle et al., 2007).

Therefore, the main objective of our study was to analyze the impact of different pure and mixed stand types (pure European beech, pure Norway spruce, pure Douglas-fir, mixed European beech/Norway spruce, mixed European beech/Douglas-fir) on nutrient conditions along Northern Germany. We studied how species identities may have shaped the nutrient status in the O-horizon and in the upper mineral soil. We hypothesized that i) the admixture of the non-native conifer species Douglas fir to beech forests increased nutrient availability, i.e. increased nutrient and C accumulation in the mineral soil, ii) in monocultures of Douglas fir and European beech the nutrient pool is comparable, but differ from pure Norway spruce stands revealing a conifer species identity effect, and iii) on nutrient poor soils species-identity effects are stronger than on rich soils.

## 2. Material and Methods

### 2.1. Study sites

We investigated seven sites in two distinct regions of Lower Saxony, Germany. The sites were grouped according to soil parent material, therefore two contrasting sites conditions were identified (Table 1).

**Table 1.**
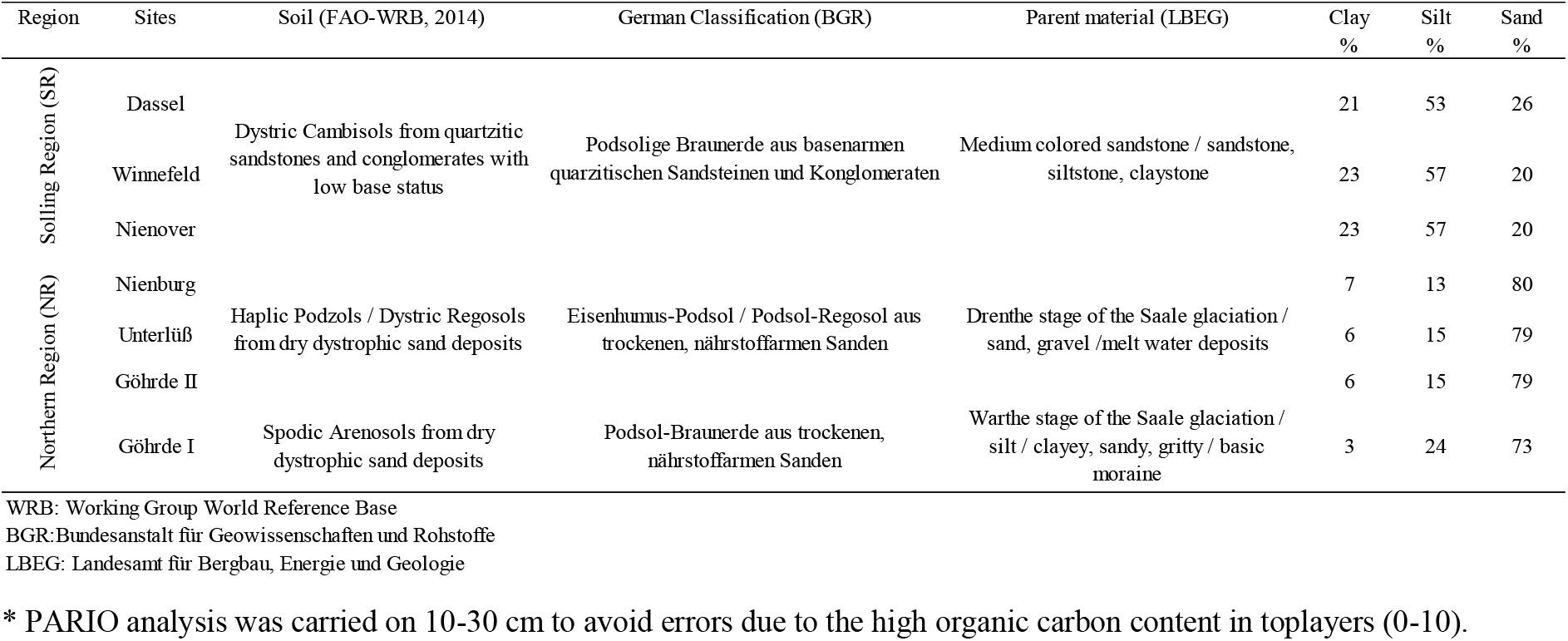
Soil classification from each site is given following (FAO, 2014) and the German classification (BGR) (Düwel et al., 2007). Soil parent material was identified matching the site coordinates with the German National inventory database (LBEG). Soil texture (10-30 cm*) was measured by the integral suspension pressure method (ISP) and determined by PARIO (see method section).

Three sites (Dassel, Winnefeld and Nienover) were chosen in South-Lower-Saxony, next to the Solling Region (SR) and are characterized by loess-influenced and rather loamy soil conditions. All sites are located between 300 and 450 m a.s.l. and are characterized by the soil type Dystric Cambisol (FAO-WRB 2015) that has developed from Triassic sandstone material (Table 1). Furthermore, all sites of the SR have similar climatic condition, i.e. a mean annual air temperature of 7.2°C and a mean annual precipitation about 1040 mm/year. The forest stand age ranged between 45 and 90 years (Table 2).

**Table 2.**
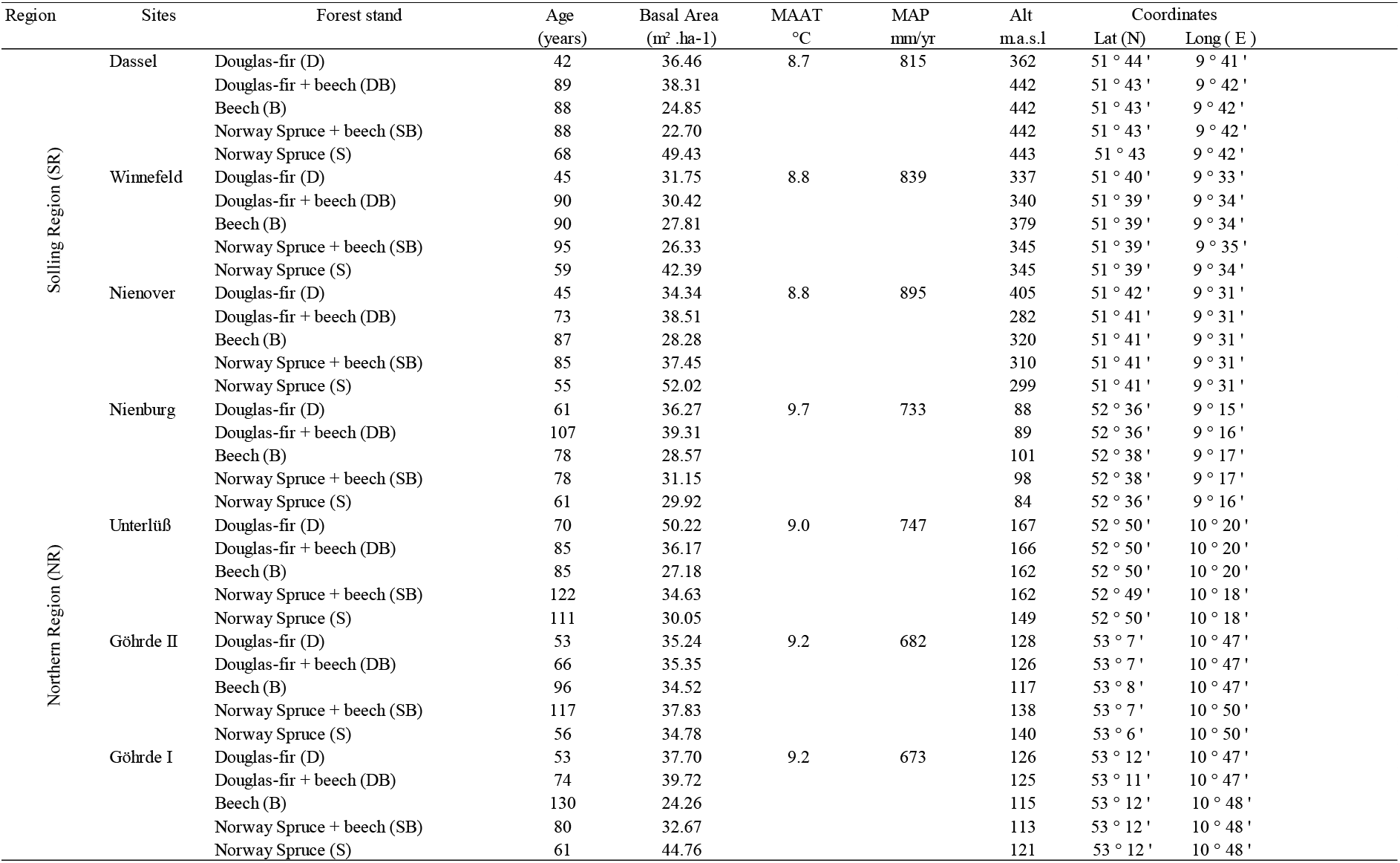
Site characteristics of each forest stand (age in years; basal area in m^2^ ha^-1^). Climate attributes were collected from climate stations of the German National Weather Service nearby each site [MAAT: Mean annual temperature; MAP: Mean annual precipitation (average from the last 20 years); ALT: Altitude]. The coordinates were taken at the center of each stand.

Additionally, four sites under sandy soil conditions were selected in the Northern part of Lower Saxony, the Northern Region (NR sites: Unterlüß, Nienburg and Göhrde I and II). The elevation ranges between 80 to 150 m a.s.l. and the soil type is characterized by a Haplic Podzol (Unterlüß, Nienburg and Göhrde II) and Spodic Arenosol (Göhrde I; FAO-WRB 2015), both soil types have developed from dystrophic sand deposits during glacial periods (Table 1). The climate condition is almost similar for all sites, i.e., a mean annual air temperature of 8.4°C, a mean annual precipitation of 720 mm/year and the forest stand age ranged between 53 and 130 years (Table 2).

At each site, five forest stands with different forest types were selected in close distance from each other (B: European beech [*Fagus sylvatica*], S: Norway spruce [*Picea abies*], D: Douglas-fir [*Pseudotsuga menziesii*] and the conifer-beech mixtures, SB: Norway spruce/Beech, DB: Douglas-fir/Beech) and within each forest stand, the distance ranged from 76 m to about 4 km (Table 2).

Liming is a common silviculture activity in Lower Saxony since the analysis of high acid inputs from atmospheric deposition since the 1970/80s. The liming and partly also fertilization activities for our research sites were accessed by German Forest Research Institute in Lower Saxony and are described in detail at the Table S1. Briefly, liming was applied on all sites in Southern area and ranged from 1 to 3 Mg ha^-1^ (also named, compensation liming, i.e. by helicopter or ground-based blower surface applied dolomite, to buffer atmospheric acid loads for 3 −5 years). The last application took place in 2010 (Solling sites). At the Northern sites, additionally to liming (ranged from 1 to 3 Mg ha^-1^), phosphate (P_2_O_5_) was applied (0.09 to 0.47 Mg ha^-1^) and the last application recorded is from 1990.

### 2.2. Soil sampling

In each pure and mixed forest stand (50 m x 50 m) at all sites, 4 randomly selected points were chosen as representative sampling points. The selected points were oriented at stand-level, e.g., we standardized two meters minimal distance from the trees to avoid coarse roots. At each sampling plot, the forest floor was collected using a steel frame and sorted by identifiable foliar (L – Litter), non-foliar (F – decay layer) and non-identifiable and humified (H – Humus) layers of the organic layer. Mineral soil was sampled using a core auger (*d*=8 cm) and, separated at 0-5, 5-10 and 10-30 cm soil depth. Bulk soil density from each depth was calculated using soil metal rings (250 cm^3^) to further stock analysis.

Partly missing bulk density data due to frozen soil conditions, interfering tree roots or stones during sampling were estimated by Adams equation (ADAMS, 1973) adapted by CHEN et al. (2017). The approach uses SOM and pH as bulk density predictors.

### 2.3. Sample preparation and analysis

All mineral soil samples were oven-dried at 40°C until constant weight and sieved through a 2 mm mesh for chemical analyses. For C and N analysis, subsamples from the fine soil fractions (d <2 mm) were grounded with a Retsch mortar grinder RM 200 (Retsch, Germany) for 10 min. Organic layer samples were dried at 60°C until constant weight, weighted and ball milled (MM2, Fa Retsch) for further chemical analyses. For soil pH analysis, 50 ml of 1 M KCl was added to 20 g of mineral soil sieved subsamples. After sedimentation of the solid phase, the pH value of the solution was determined with a glass electrode. As all mineral soil samples were negatively tested for free carbonates (HCL-test), the NH_4_Cl-percolation method according to Höhle et al. (2018) was used to determine exchangeable cation (Ca^2+^, Mg^2+^, Na^+^, K^+^ Al^3+^, Fe^2+^, Mn^2+^, H^+^) concentration. Briefly, 2.5 g sieved mineral soil was extracted by shaking the samples with 100 ml of 0.5 M NH_4_Cl solution for 2 hours. The suspension was left standing for another 24 h and afterwards filtrated through membrane filters with mesh size 0.45 μm (Sartorius, Göttingen, Germany). The cation concentrations of the filtrates were analyzed by ICP-OES (Spectro Genesis, Spectro, Kleve, Germany). Exchangeable H^+^ were calculated according to Meiwes et al., (1986) considering the given pH and the aluminum concentration in the percolate. The sum of all extracted cations was defined as the effective cation exchange capacity (CEC; mmol_c_ kg^-1^). The base saturation (BS) was defined as the share of exchangeable cations Ca^2+^, Mg^2+^, K^+^ and Na^+^ on the CEC.

The concentration of Ca, K, Mg and P from organic layers were determined by pressure digestion with 65 % nitric acid for 8 h at 170°C (Höhle et al., 2018). Digestates were filtered by ash-free cellulose filters and determined by ICP-OES (Spectro Genesis, Spectro, Kleve, Germany). Further details on the applied analytical procedure can be also found in König et al., (2014).

We estimated the soil bulk density from the oven-dried and moisture corrected (105 °C) fine soil mass and its volume. The fine soil volume was estimated from the difference between the volume of the soil corer and the volume of stones and roots. Organic layer nutrients stocks were calculated multiplying nutrient concentration by organic layer mass. Nutrients stocks in each mineral soil layer were calculated from the soil bulk density, concentrations of nutrient and depth of the soil layer.

Soil texture was measured by integral suspension pressure method (ISP) (Durner et al., 2017) and determined by PARIO (METER Group, Inc. USA).

### 2.4. Statistical Analyses

To address the non-independent nature of multiple horizons within one soil profile for this subset of data, linear mixed effect models were chosen for this statistical approach as in applied by Rasmussen et al. (2018). To estimate the effect of forest stand and site condition (loamy vs sandy), we fitted linear mixed models (LMMs) to log-transformed response variables (CEC, pH, exchangeable cations stocks and organic layer nutrient stocks [Ca, Mg and K]) and then applied planned contrasts (Piovia-Scott et al., 2019). All LMMs included forest stands (European beech, Douglas-fir, Douglas-fir/European beech, Norway spruce, Norway spruce/European beech), site conditions (loamy and sandy sites) and soil depths (Ol, Of, Oh [organic layers] and 0-5, 5–10 and 10–30 cm [mineral soil]) as fixed effects. Models were stepwise selected by likelihood ratio test, and minimal models included all main effects and the interaction of forest type and site texture.

Additionally, in the complementary material the mean values for each site are presented (Table S2 and S3). There, the effect of forest stand (pure or mixed forest) on the soil chemical parameters for individual depth was assessed by LSD test. As in some cases the assumption of normal distribution was not met (tested for with Shapiro–Wilk–test), the Kruskal–Wallis–H–test, followed by pairwise Mann–Whitney–U–tests with a correction factor for multiple pairwise testing, was used to identify statistically significant differences among forest stand and sites conditions, by individual depth.

All analyses were done in R 4.0.3 (R Core Team, 2020). We used the ‘nlme’ package to fit LMMs and the ‘emmeans’ package to conduct planned contrasts. All mixed models met the assumptions of normality of residuals and homogeneity of variance.

## 3. Results

### 3.1. Soil base saturation, pH and exchangeable cations

The base saturation of the mineral topsoil (0-5 cm) ranges from 30% on B and D stands, to minimum values around 18% on its mixtures (DB). Intermediate values were observed under S stands and its mixtures (SB), 25%. For the soil depth 10-30 cm, the majority of the measured base saturation values fall below 10%, with minimum values of 7% in SB stands and up to maximum values of 12% on pure D and B stands.

Significant tree species and species mixtures impact on soil base saturation (BS) occurred only at the Solling sites (SR) (5-10 cm; Table S2). There, pure B stands showed similar base saturation compared to mixed stands, ranging from 9 to 14% and lower values than both conifers, with 19 % for D and 15% for S stands. The base saturation (0-5 cm) was 22% higher under the mixed Douglas-beech (DB) than SB stands (NR sites). The opposite was observed at the SR sites, where the base saturation was 27% lower under DB compared to SB stand, but nonstatistical differences were observed.

The cations exchangeable capacity (CEC) of the mineral topsoil (0-5) ranges from 124 mmol_c_ kg^-1^ on the SR sites to 54.3 mmol_c_ kg^-1^ on NR sites, both values were observed under S stands (Figure 1a). Intermediate CEC ranges from 105 mmol_c_ kg^-1^ at SR sites under B, D and the mixtures (SB and DB) to 73 mmol_c_ kg^-1^ at NR sites under DB. For the soil depth (10-30) the CEC falls below 50 mmol_c_ kg^-1^, with maximum CEC of 58 mmol_c_ kg^-1^ under DB (SR) and minimum of 23 mmol_c_ kg^-1^ under B at NR sites.

**Figure 1.**
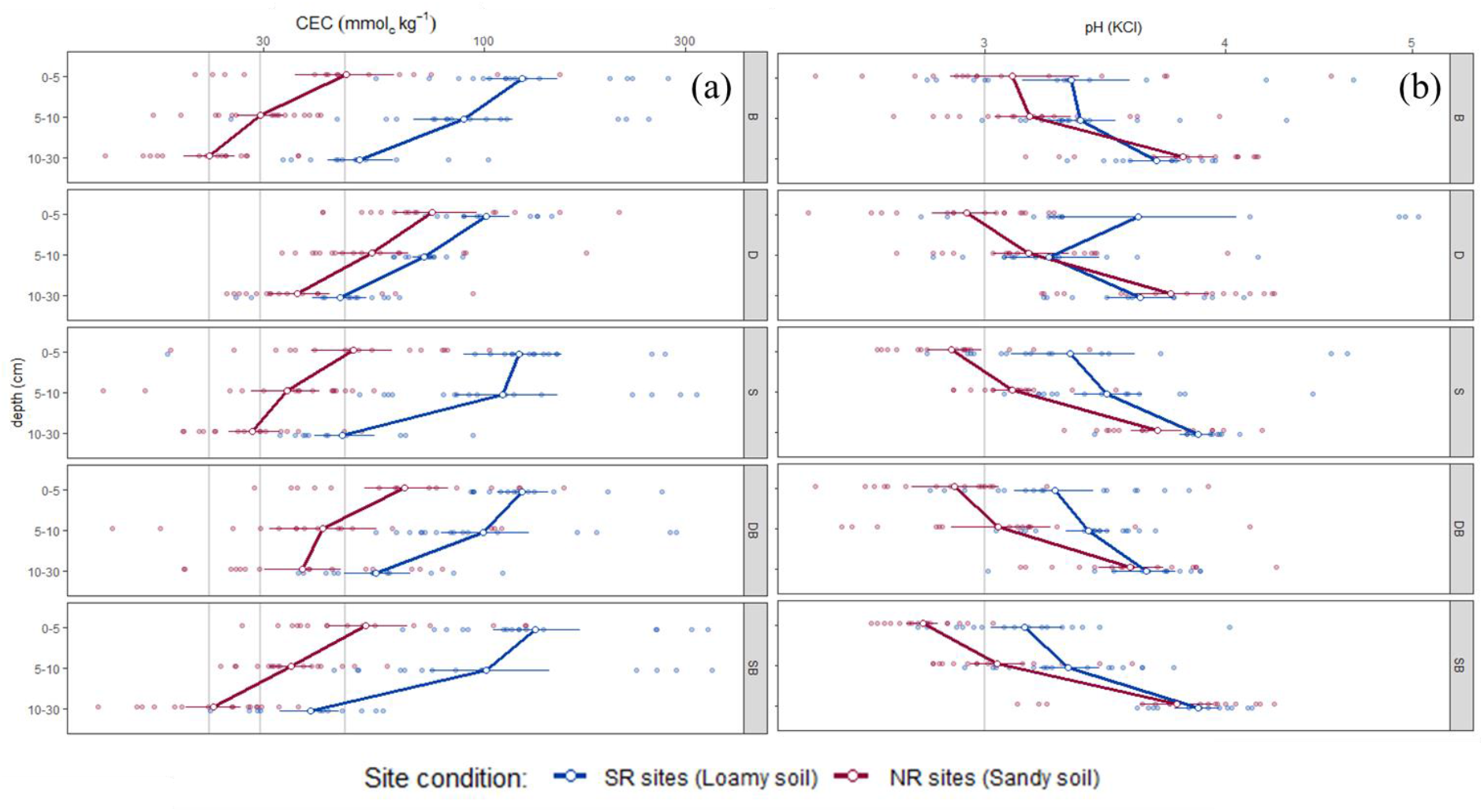
Changes in CEC (a) (mmol_c_ kg^-1^) and pH (b) (KCl) with mineral soil depth (0-5, 5-10 and 10-30 cm) and forest stands (B: European beech, D: Douglas-fir, S: Norway spruce and the conifer-beech mixtures, SB: Norway spruce/Beech, DB: Douglas-fir/Beech) at SR (loamy soil; *n*=12) and NR (Sandy soil; *n*=16). White point represents the mean and horizontal bars the standard errors.

The results of the linear mixed effect model, testing the interactions between forest stand (F), site condition (S) and soil horizon (D) (Table 3), showed a significant influence of the interaction between forest stand and site condition (P<0.05; Table 3) and depth and site condition (P< 0.01; Table 3). The forest stand effects were significant at the NR sites, but not in SR sites. The variation of CEC among forest under sandy soil conditions was higher under pure D stands (84.0 mmol_c_.kg^1^) compared to pure S (54.3 mmol_c_.kg^1^) and B (55.3 mmol_c_.kg^1^) stands (Table S2; Figure 1a).

**Table 3.** The F- and P-values of linear mixed-effects models on the effect of Forest stand (B: European beech, D: Douglas-fir, S: Norway spruce and the conifer-beech mixtures, SB: Norway spruce/Beech, DB: Douglas-fir/Beech), Site condition (SR -loamy and sandy sites) and depth (0-5, 5-10 and 10-30 cm) on CEC (mmol_c_ kg^-1^), pH (KCl) and Base saturation (%) (a), exchangeable cation (Ca ^2+^, Mg^2+^ and K ^+^; mmol_c_ kg^-1^) (b) and, exchangeable cation stocks (Ca ^2+^, Mg^2+^ and K^+^; kg ha^-1^). Significant effects are given in bold (p < 0.05). For mean values please refer to the Table S2.

| (a) | CEC ($\text{mmol}_c \text{kg}^{-1}$) | | | pH (KCl) | | | BS (%) | | |
| --- | --- | --- | --- | --- | --- | --- | --- | --- | --- |
|  | df | F | P | df | F | P | df | F | P |
| Forest stand (F) | 23 | 2.84 | **0.05** | 23 | 0.42 | 0.79 | 23 | 1.55 | 0.22 |
| Depth (D) | 406 | 153.28 | **<.0001** | 406 | 139.63 | **<.0001** | 406 | 125.63 | **<.0001** |
| Site condition (S) | 6 | 12.45 | **0.01** | 6 | 8.02 | **0.03** | 6 | 0.01 | 0.93 |
| F x S | 23 | 2.64 | **0.05** | 23 | 0.13 | 0.97 | 23 | 0.3 | 0.87 |
| D x S | 406 | 4.43 | **0.01** | 406 | 11.7 | **<.0001** | 406 | 3.49 | **0.03** |

| (b) | $\text{Ca}^{2+}$ ($\text{mmol}_c \text{kg}^{-1}$) | | | $\text{Mg}^{2+}$ ($\text{mmol}_c \text{kg}^{-1}$) | | | $\text{K}^{+}$ ($\text{mmol}_c \text{kg}^{-1}$) | | |
| --- | --- | --- | --- | --- | --- | --- | --- | --- | --- |
|  | df | F | P | df | F | P | df | F | P |
| Forest stand (F) | 23 | 2.26 | **0.09** | 23 | 3.07 | **0.04** | 23 | 2.58 | 0.06 |
| Depth (D) | 406 | 183 | **<.0001** | 406 | 231.21 | **<.0001** | 406 | 146.38 | **<.0001** |
| Site condition (S) | 6 | 3.71 | 0.1 | 6 | 8.86 | **0.02** | 6 | 29.55 | **<.0001** |
| F x S | 23 | 0.1 | 0.98 | 23 | 0.6 | 0.67 | 23 | 1.42 | 0.26 |
| D x S | 406 | 1.9 | 0.15 | 406 | 3.31 | **0.04** | 406 | 6.67 | **<.0001** |

| (c) | Exchangeable Ca stock ($\text{Kg ha}^{-1}$) | | | Exchangeable Mg stock ($\text{Kg ha}^{-1}$) | | | Exchangeable K stock ($\text{Kg ha}^{-1}$) | | |
| --- | --- | --- | --- | --- | --- | --- | --- | --- | --- |
|  | df | F | P | df | F | P | df | F | P |
| Forest stand (F) | 23 | 1.96 | 0.13 | 23 | 2.89 | **0.04** | 23 | 1.91 | 0.14 |
| Depth (D) | 406 | 34.64 | **<.0001** | 406 | 43.07 | **<.0001** | 406 | 383.03 | **<.0001** |
| Site condition (S) | 6 | 3.42 | 0.11 | 6 | 8.3 | **0.03** | 6 | 29.88 | **<.0001** |
| F x S | 23 | 0.19 | 0.94 | 23 | 0.66 | 0.62 | 23 | 1.41 | 0.26 |
| D x S | 406 | 1.86 | 0.16 | 406 | 3.37 | **0.04** | 406 | 12.94 | **<.0001** |

The pH values were highest under D stands at SR sites (3.7), meanwhile the lowest pH was observed under SB stands (2.7) at NR sites. No interaction between forest stand and site condition were observed (P<0.97; Table 3). However, a tendency was observed at NR sites, where B showed the highest pH (3.1) and S, DB and SB the lowest (~2.8), D stands presented intermediate results (2.9) (Figure 1b). At the SR sites, the pH values below 10 cm differed significantly between the pure stands (S, B, D). At this depth, the pH ranged from 3.8 under S stand to 3.6 under B and D stands. Independent of the sites, the pH values under mixture stands were mainly between those of the respective pure stands (Figure 1b), except under SB stands at NR sites, where the lowest pH was observed. The linear mixed effects showed a significant influence of site conditions on pH (P<0.03), where the mean pH at SR sites were higher than at the NR sites.

Mineral topsoil exchangeable Al concentrations was higher under pure D and S (30.9 and 18.9 mmol_c_kg^1^, respectively) compared to B stand (13.31 mmol_c_kg^1^) at NR sites, meanwhile at SR sites, B and S showed no significance differences on Al concentrations (43.9 and 60.72 mmol_c_·kg^1^, respectively) but higher than D stands (20.89 mmol_c_kg^1^). Forest stand effects on deeper mineral soil (10–30) were small and inconsistent at both regions (Table S2).

The C:N ratio of mineral topsoil (0-5) ranges from 18 under conifers (D and S) and both mixed stands (DB and SB) to 16 under B stand at the SR sites. Even though higher values were observed at NR sites (ranging from 25 to 19), the same pattern was observed. Minimum values were observed under B stands (19) and maximum values (~24) under D, S and both mixed stands (DB and SB). At the depth 10-30 cm, the C:N ratio values were below 15 (SR sites) and below 24 (NR sites). Significant differences were observed only at the NR sites, maximum value of 23 under D stands and minimum value of 17 under B stand, intermediate values were observed under S stands (20) and SB stands (18).

### 3.2. Exchangeable base cations stocks in the upper mineral soil

Patterns of soil exchangeable base cations stocks differed depending on site, forest stand and depth (Table 3; Figure 2). At SR sites, the mineral topsoil (0-5) exchangeable Ca stocks ranges from 37 kg ha^-1^ under D stands to about 10 kg ha^-1^ under the SB mixed stands. At NR sites, Ca stocks in the 0-5 cm soil depth are reduced, compared to the SR, reaching highest values under B and D (15 and 12 kg.ha^-1^, respectively) and lowest under SB. In the 5-10 but also in the 10 to 30 cm soil depths, Ca stocks were even stronger reduced to about 10 or below 5 kg ha^-1^ (Figure 2).

**Figure 2.**
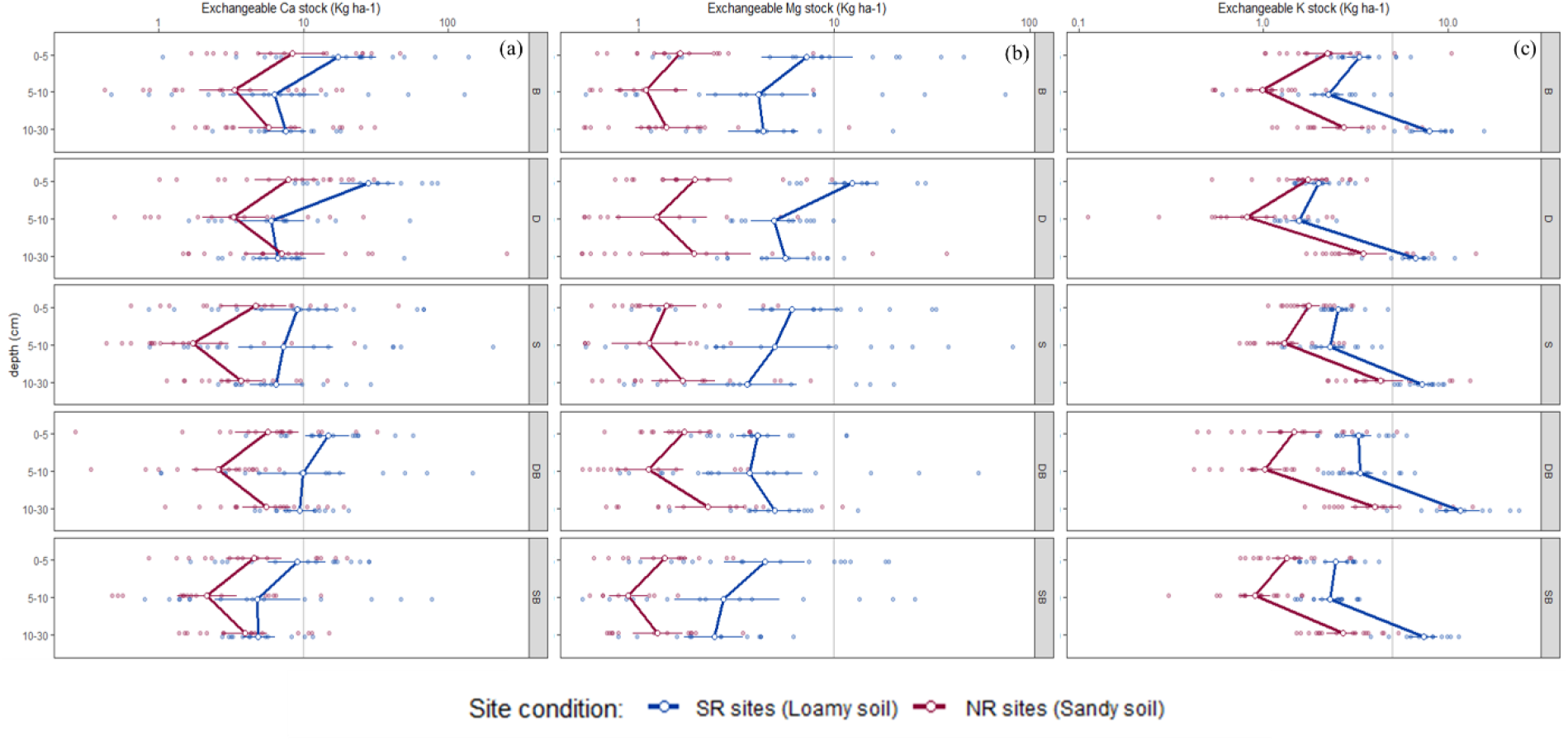
Changes of exchangeable base cations stocks Ca (a), Mg (b) and K (c) (kg ha^-1^) at different mineral soil depth (0-5, 5-10 and 10-30 cm) and forest stands (B: European beech, D: Douglas-fir, S: Norway spruce and the conifer-beech mixtures, SB: Norway spruce/Beech, DB: Douglas-fir/Beech) at SR (loamy soil; *n*=12) and NR (Sandy soil; *n*=16). White point represents the mean and horizontal bars the standard errors.

The same pattern was observed for exchangeable Mg stocks. These values range from 13 kg ha^-1^ under D stands to below 5 kg ha^-1^ under mixtures stands (DB and SB) at SR sites (0-5 cm). At the NR, stocks of Mg often showed values below 2.5 kg ha^-1^ under all forests and depth, expect under D stands (10-30 cm), where the stocks of 5 kg ha^-1^ were observed.

The exchangeable K stocks ranged from 12 kg ha ^-1^ under DB at the SR sites to below 2 at the NR sites. At the SR sites, the mixture DB showed the highest K stocks (all layers) followed by pure B. Both conifers (D and S) showed the lowest K stocks, expect at the NR sites (10-30), where the K stock was comparable to the DB stand.

### 3.3. Nutrient stocks in the organic layer

Nutrient stocks in the organic layers differed between tree species stands and sites. For all layers, high Mg and K stocks were observed at the SR sites, meanwhile low stocks were found at the NR sites (Figure 3). No differences between region were observed for Ca stocks. In general, higher Ca, Mg and K stocks were observed under both conifers (D and S) than under B stands, but the pattern differs among sites. At the SR sites, D stands stored 53.6 kg ha^-1^ of Ca (L-layer) and 86.9 kg ha^-1^ (H-layer), meanwhile S stands accumulated 32.5 kg ha^-1^ (L-layer) and 51.0 kg ha^-1^ (H-layer). At NR sites the opposite was observed, higher stocks of Ca were observed under S stands (47.0 kg ha^-1^ L-layer; 90.1 kg ha^-1^ H-layer) than D stands (30.2 kg ha^-1^ L-layer; 31.2 kg ha^-1^ H-layer). Meanwhile, under B stands the Ca storage at the L-layer did not differ between sites (28.7 kg ha^-1^ at SR and 29 kg ha^-1^ at NR), at the H-layer higher stocks of Ca was observed at SR (39.1 kg ha^-1^) than at NR sites (26.7 kg ha^-1^).

**Figure 3.**
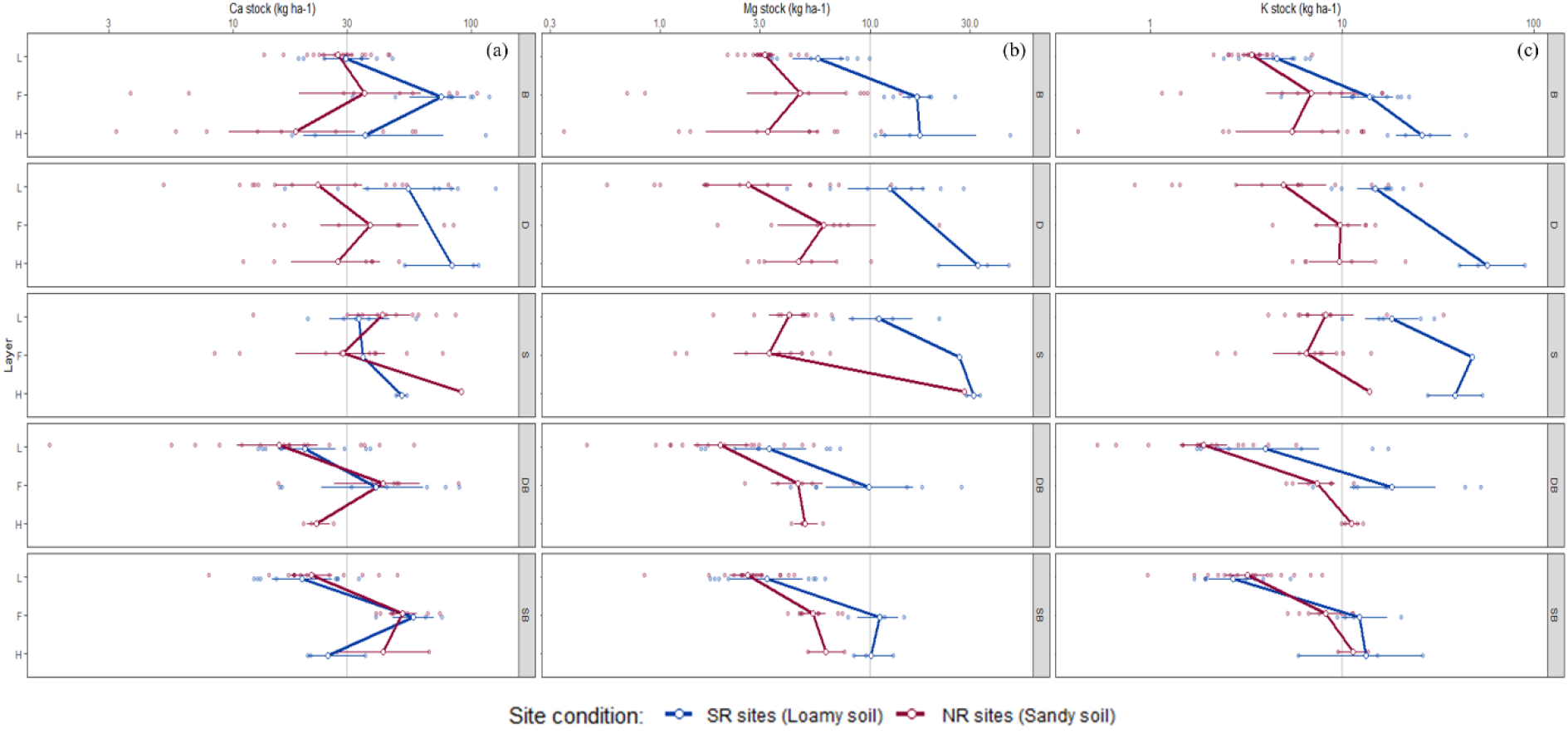
Changes in Ca (a), Mg (b) and K (c) stocks (kg ha^-1^) at organic layers (L, F and H) and forest stands (B: European beech, D: Douglas-fir, S: Norway spruce and the conifer-beech mixtures, SB: Norway spruce/Beech, DB: Douglas-fir/Beech) at SR (loamy soil; *n*=12) and NR (Sandy soil; *n*=16). White point represents the mean and horizontal bars the standard errors.

All species showed higher storages of Mg at the SR sites than at the NR sites. At both regions, higher Mg stocks at the L-layer were observed under D (12.0 kg ha^-1^ SR; 3.9 kg ha^-1^ NR) and S (10.8 kg ha^-1^ SR; 4.3 kg ha^-1^ NR) stands, compared to B (5.1 kg ha^-1^ SR; 3.2 kg ha^-1^ NR) stands. The highest storage was observed at the H-layer, but significant differences were observed only at the NR sites (P< 0.01), where Mg storage under S stands was 80 % higher than D stands.

Similar than observed for Mg, the K storage were significantly higher at the SR sites than at the NR sites (P<0.006) and both conifers showed significantly higher K stocks at the L-layer, compared to B in both regions. At the L-layer, higher K storage was observed under S (21.9 kg ha^-1^ SR; 10.0 kg ha^-1^ NR) stands than D stand (14.2 kg ha^-1^ SR; 7.6 kg ha^-1^ NR), meanwhile at the H-layer the opposite was observed at the SR sites. There, the higher stocks were observed under D (60.2 kg ha^-1^ SR; 11.0 kg ha^-1^ NR) than S (40.6 kg ha^-1^ SR; 13.9 kg ha^-1^ NR).

The mixed stands often showed intermediate storage between the conifers and beech for all nutrients analyzed (Ca, Mg and K). The only difference between the mixtures were observed at the NR sites, where the storages of K under SB stands were significantly higher than DB (L-layer).

### 3.4 Vertical distribution and sum of element stocks in the upper soil horizons

The vertical distributions and the sums of the respective stocks of Ca, Mg and K, from the organic layer and the mineral topsoil for all sites and forest stand are summarized at Table 4. First, it is striking that only with the exception of Ca and the stocks for the S, DB and SB stands, all other results indicate a clearly increased stock of nutrient at the SR sites.

**Table 4.** The F- and P-values of linear mixed-effects models on the effect of Forest stand (European beech, Douglas-fir, Norway spruce, mixture of European beech with Douglas-fir and mixture of European beech with Norway spruce), Site condition (loamy and sandy sites) and depth (L, F and H layer) on Ca, Mg and K stocks (kg ha^-1^). Significant effects are given in bold (p < 0.05). For mean values for each site/forest stand, refer to Table S3.

| | Ca ($\text{kg ha}^{-1}$) | | | Mg ($\text{kg ha}^{-1}$) | | | K ($\text{kg ha}^{-1}$) | | |
| --- | --- | --- | --- | --- | --- | --- | --- | --- | --- |
|  | df | F | P | df | F | df | DF | F | P |
| Forest stand (F) | 16 | 3.82 | **0.022** | 16 | 3.6 | **0.02** | 16 | 6.95 | **<.0001** |
| Depth (D) | 171 | 14.53 | **<.0001** | 171 | 37.09 | **<.0001** | 171 | 61.11 | **<.0001** |
| Site condition (S) | 4 | 17.28 | **0.014** | 4 | 25.46 | **0.007** | 4 | 27.33 | **0.006** |
| F x S | 16 | 6.98 | **<.0001** | 16 | 2.61 | 0.074 | 16 | 2.03 | 0.138 |
| D x S | 171 | 3.98 | **0.02** | 171 | 4.63 | **0.01** | 171 | 4.68 | **0.01** |

**Table 4.** Stocks of Ca, Mg and K (kg ha^-1^) of individual soil profile (Organic layer: L, F and H; Mineral soil (MS): 0-30 cm) (mean value ± standard error) under different forest stand (B = European beech, D = Douglas fir, S = Norway spruce, DB = Douglas fir-beech mixture and SB = spruce-beech mixture) at Solling region sites (SR) and Northern region sites (NR). For the statistical analysis refer to Table S1 and S2.

| Site | Species | depth | Ca stock (kg ha<sup>-1</sup>) | Mg stock (kg ha<sup>-1</sup>) | K stock (kg ha<sup>-1</sup>) | Site | Species | depth | Ca stock (kg ha<sup>-1</sup>) | Mg stock (kg ha<sup>-1</sup>) | K stock (kg ha<sup>-1</sup>) |
| --- | --- | --- | --- | --- | --- | --- | --- | --- | --- | --- | --- |
| Solling | B | L | 30.9 ± 3.5 | 6.2 ± 0.9 | 4.8 ± 0.6 | Northern | B | L | 29.0 ± 2.3 | 3.2 ± 0.2 | 3.5 ± 0.3 |
|  |  | F | 80.4 ± 10.1 | 17.2 ± 1.5 | 15.1 ± 2.1 |  |  | F | 50.3 ± 9.9 | 6.3 ± 1.2 | 9.0 ± 1.6 |
|  |  | H | 47.9 ± 22.7 | 21.1 ± 8.6 | 27.8 ± 5.9 |  |  | H | 26.7 ± 6.6 | 4.7 ± 1.1 | 7.7 ± 1.5 |
|  |  | Total Organic layer | 159.2 | 44.5 | 47.8 |  |  | Total Organic layer | 106.0 | 14.2 | 20.2 |
|  |  | 0-5 | 17.8 ± 4.1 | 6.5 ± 1.6 | 3.2 ± 0.3 |  |  | 0-5 | 13.9 ± 3.5 | 1.9 ± 0.5 | 2.8 ± 0.6 |
|  |  | 5-10 | 4.8 ± 1.0 | 3.0 ± 0.6 | 2.1 ± 0.2 |  |  | 5-10 | 5.9 ± 1.5 | 1.1 ± 0.5 | 1.1 ± 0.1 |
|  |  | 10-30 | 8.8 ± 1.4 | 5.7 ± 1.4 | 8.5 ± 0.9 |  |  | 10-30 | 9.4 ± 2.3 | 2.0 ± 0.7 | 3.2 ± 0.5 |
|  |  | Total MS | 31.4 | 15.2 | 13.8 |  |  | Total MS | 29.2 | 5.1 | 7.1 |
|  |  | TOTAL | 190.5 | 59.7 | 61.6 |  |  | TOTAL | 135.2 | 19.3 | 27.3 |
| Solling | D | L | 64.9 ± 12.9 | 14.6 ± 2.8 | 15.4 ± 1.5 | Northern | D | L | 30.2 ± 6.4 | 3.8 ± 1.0 | 7.6 ± 2.0 |
|  |  | F | NA |  |  |  |  | F | 45.5 ± 10.5 | 7.7 ± 2.4 | 10.4 ± 1.4 |
|  |  | H | 86.9 ± 17.4 | 34.4 ± 7.2 | 60.3 ± 14.9 |  |  | H | 31.3 ± 6.2 | 5.2 ± 1.2 | 11.0 ± 2.5 |
|  |  | Total Organic layer | 151.8 | 48.9 | 75.7 |  |  | Total Organic layer | 107.0 | 16.7 | 29.0 |
|  |  | 0-5 | 37.0 ± 7.7 | 13.9 ± 2.1 | 2.1 ± 0.2 |  |  | 0-5 | 11.5 ± 2.1 | 2.6 ± 0.7 | 1.9 ± 0.2 |
|  |  | 5-10 | 10.3 ± 4.2 | 5.5 ± 0.7 | 1.6 ± 0.1 |  |  | 5-10 | 5.7 ± 1.7 | 1.4 ± 0.5 | 1.0 ± 0.2 |
|  |  | 10-30 | 9.7 ± 3.7 | 6.3 ± 0.8 | 6.9 ± 0.5 |  |  | 10-30 | 24.2 ± 15.4 | 4.9 ± 2.4 | 4.3 ± 0.8 |
|  |  | Total MS | 57.0 | 25.8 | 10.5 |  |  | Total MS | 41.4 | 9.0 | 7.2 |
|  |  | TOTAL | 208.8 | 74.7 | 86.2 |  |  | TOTAL | 148.4 | 25.7 | 36.2 |
| Solling | S | L | 35.6 ± 6.3 | 12.0 ± 2.6 | 19.5 ± 3.7 | Northern | S | L | 46.8 ± 6.2 | 4.3 ± 0.4 | 10.0 ± 2.6 |
|  |  | F | 34.9 ± | 26.6 ± | 47.7 ± |  |  | F | 35.1 ± 7.0 | 3.8 ± 0.6 | 7.5 ± 1.2 |
|  |  | H | 50.8 ± 2.6 | 31.2 ± 2.4 | 40.6 ± 12.8 |  |  | H | 90.8 ± | 28.2 ± | 13.9 ± |
|  |  | Total Organic layer | 121.3 | 69.7 | 107.7 |  |  | Total Organic layer | 172.8 | 36.3 | 31.4 |
|  |  | 0-5 | 17.9 ± 6.5 | 8.5 ± 2.7 | 2.7 ± 0.2 |  |  | 0-5 | 8.5 ± 2.8 | 1.6 ± 0.3 | 1.8 ± 0.2 |
|  |  | 5-10 | 8.8 ± 4.1 | 5.2 ± 2.0 | 2.0 ± 0.1 |  |  | 5-10 | 3.5 ± 1.4 | 0.9 ± 0.2 | 1.4 ± 0.2 |
|  |  | 10-30 | 8.8 ± 2.4 | 6.0 ± 1.9 | 7.4 ± 0.5 |  |  | 10-30 | 4.8 ± 0.9 | 2.1 ± 0.5 | 5.1 ± 0.8 |
|  |  | Total MS | 35.5 | 19.6 | 12.1 |  |  | Total MS | 16.8 | 4.6 | 8.3 |
|  |  | TOTAL | 156.8 | 89.4 | 119.8 |  |  | TOTAL | 189.6 | 40.9 | 39.7 |
| Solling | DB | L | 21.9 ± 3.7 | 3.9 ± 0.8 | 6.1 ± 2.2 | Northern | DB | L | 20.7 ± 3.7 | 2.3 ± 0.3 | 2.2 ± 0.3 |
|  |  | F | 48.3 ± 11.1 | 12.1 ± 3.2 | 22.9 ± 6.7 |  |  | F | 47.8 ± 9.6 | 4.9 ± 0.8 | 7.7 ± 1.0 |
|  |  | H | NA |  |  |  |  | H | 22.5 ± 1.4 | 4.9 ± 0.4 | 11.3 ± 0.7 |
|  |  | Total Organic layer | 70.2 | 16.0 | 29.0 |  |  | Total Organic layer | 91.0 | 12.1 | 21.2 |
|  |  | 0-5 | 20.9 ± 4.9 | 4.2 ± 0.8 | 3.9 ± 0.3 |  |  | 0-5 | 8.8 ± 2.1 | 1.7 ± 0.3 | 1.9 ± 0.3 |
|  |  | 5-10 | 7.6 ± 1.7 | 2.5 ± 0.5 | 3.3 ± 0.4 |  |  | 5-10 | 3.3 ± 0.5 | 1.2 ± 0.3 | 1.1 ± 0.1 |
|  |  | 10-30 | 10.5 ± 1.4 | 5.7 ± 0.9 | 12.8 ± 1.7 |  |  | 10-30 | 7.1 ± 1.2 | 3.0 ± 0.7 | 4.8 ± 0.8 |
|  |  | Total MS | 38.9 | 12.4 | 20.0 |  |  | Total MS | 19.2 | 5.9 | 7.8 |
|  |  | TOTAL | 109.1 | 28.4 | 49.0 |  |  | TOTAL | 110.2 | 18.0 | 29.0 |
| Solling | SB | L | 21.0 ± 3.1 | 3.7 ± 0.7 | 2.9 ± 0.5 | Northern | SB | L | 23.2 ± 2.6 | 2.8 ± 0.2 | 3.6 ± 0.5 |
|  |  | F | 58.6 ± 7.4 | 11.3 ± 1.4 | 12.9 ± 2.5 |  |  | F | 52.4 ± 4.1 | 5.5 ± 0.4 | 8.5 ± 0.9 |
|  |  | H | 25.8 ± 5.0 | 10.3 ± 1.3 | 15.8 ± 5.8 |  |  | H | 46.7 ± 19.4 | 6.3 ± 1.2 | 11.6 ± 2.1 |
|  |  | Total Organic layer | 105.4 | 25.3 | 31.6 |  |  | Total Organic layer | 122.3 | 14.6 | 23.8 |
|  |  | 0-5 | 10.6 ± 2.5 | 4.9 ± 1.5 | 2.8 ± 0.2 |  |  | 0-5 | 6.8 ± 1.5 | 1.3 ± 0.2 | 1.5 ± 0.2 |
|  |  | 5-10 | 3.2 ± 0.8 | 1.6 ± 0.3 | 2.2 ± 0.2 |  |  | 5-10 | 3.3 ± 0.8 | 0.6 ± 0.1 | 1.0 ± 0.1 |
|  |  | 10-30 | 5.5 ± 0.9 | 2.9 ± 0.5 | 7.7 ± 0.7 |  |  | 10-30 | 5.2 ± 1.0 | 1.2 ± 0.2 | 2.9 ± 0.3 |
|  |  | Total MS | 19.2 | 9.3 | 12.7 |  |  | Total MS | 15.3 | 3.1 | 5.4 |
|  |  | TOTAL | 124.6 | 34.7 | 44.2 |  |  | TOTAL | 137.6 | 17.6 | 29.1 |

Overall, Ca stocks in both regions are in the range of about 110 to about 200 kg ha^-1^ and are significantly increased only under B and D stands at the SR sites, compared to the NR sites. Noticeable elevated (S stand) in the NR sites and almost comparable Ca stocks were found under DB and SB stands in both regions. With reference to the vertical distribution in the topsoil, it is striking that only between about 15 to 35% (SR) and about 9 to 28% (NR) of the considered upper topsoil Ca stocks are bound in the mineral soil. It is still apparent that higher Ca stocks in the upper mineral are mostly found with the participation of B, D or their mixture (DB) in both regions.

The total stocks of Mg at the SR sites range from about 28 - 90 kg ha^-1^, with highest values under S and D, to medium values under B, and to lowest values under mixtures, DB and SB. The percentage in the upper mineral soil of the total determined topsoil stock ranges from just over 20% under S to over 40% under DB. At the NR, the total Mg stocks range from about 18 to 40 kg ha^-1^. Here, the comparatively lowest mineral soil percentages of the total stock were found under S (11%), and the relatively highest under D (35%). Furthermore, it is clear that in both study regions, the highest total stocks of Mg were determined under S and the lowest in the two mixed stands DB and SB.

Total K stocks at the SR sites ranges from 44 to 120 kg ha^-1^, with high values under S, lower to medium values under D and B, to relatively lowest values under mixed stands, DB and SB. The percentage in the upper mineral soil of the total K stocks ranges from 10% under S to over 40% under DB. At NR sites, total K stocks range from 30 to 40 kg ha^-1^. Here, the comparatively highest percentage of K in the mineral soil was found under pure B (26%) or its mixture with Douglas-fir (DB) (27%). The relatively lowest percent of K stocks in the mineral soil with about 20% were analyzed for the pure stands of D and S as well as for the mixture SB.

## 4. Discussion

### Context and approach of the work

The toplayer of the forest soil is probably the most important soil compartment with regard to the turnover of nutrients and organic matter (Binkley, 2019; Binkley and Giardina, 1998), e.g. through leaf litter, intensive rooting, atmospheric input via throughfall, as well as the accumulation of organic matter and the associated soil biological activities. Thereby, the example of neighboring comparison stands from the dataset of the soil condition survey BZE (Bodenzustandserhebung) in Lower Saxony (Evers et al., 2019) shows significant depth gradients with respect to the preferential accumulation of nutrient element cations (Ca, Mg, K) in the organic layer and a clear depletion in the upper mineral soil up to a soil depth of 30 cm, especially among old spruce stands on the sandy and thus also tendentially nutrient-poorer sites in the northern region (example site BZE NI00504). The same is true for spruce in Solling on topsoil acidified Triassic sandstone material (example site BZE NI00047). Also under beech, comparative stands from the BZE data set show a clear depth gradient under the corresponding soil texture in both study regions (example sites BZE NI00018, Solling Region; BZE NI00695, Northern Region). However, here the level of accumulated nutrient cations in the total mineral soil is often higher, which is usually due to the fact that in deeper soil horizons a connection by the tree roots to less acidified or to richer parent substrates (e.g. loams or boulder sands) is given.

Thus, it remains to be stated that the reduction of the considerations made in the present study to soil chemical characteristic data for the organic layers down to a mineral soil depth of 30 cm is not suitable for providing a comprehensive overall assessment of the respective soil chemical site conditions. At the same time, however, on the background of existing data sets from e.g., the comprehensive soil condition survey in the forest or from specific studies on rooting (Lwila et al., 2021) or soil biology (Lu and Scheu, 2020), it can be deduced that the topsoil investigated here forms an essential interface to the soil biology and to the growing stock solely by intensive root growth or litter input. It was therefore assumed that not only the largely known and tree species-specific properties of beech and spruce (humus form, accumulation of litter, rooting pattern; Augusto et al., 2002; Konôpka, 2009; Grüneberg et al., 2019), but also, those of Douglas-fir (tending to lower litter C:N ratios, compared to spruce; Trum et al., 2011) and their mixture with beech, compared to the mixture with spruce can be mapped via the soil chemical characterization of the forest topsoil. In addition to the site-comparative consideration of the standard soil parameters, the main focus is on the nutrient element cations Ca, Mg and K. The reasons for this are that these elements often fall into critical ranges on poorer parent substrates and under the generally prevailing topsoil acidification due to increased atmospheric acid deposition (Meesenburg et al., 2019). Conversely, according to the studies available so far and related to the production of woody biomass, a high utilization efficiency is indicated for the Douglas fir of these elements (Block et al., 2008; Ranger et al., 1995).

### Topsoil acidification among beech and conifers are site dependent

Significant differences in soil acidification were observed between regions. The SR sites showed less soil acidification (pH ~ 3.8) and a significantly enhanced CEC (~100 mmol_c_ kg^-1^ at 0-5 cm), compared to the NR sites (pH: ~ 3; CEC: 60 mmol_c_ kg^-1^ at 0-5 cm). At the SR, S stands showed less acidic conditions than D and B, whereby the opposite was observed at the NR sites, where a greater soil acidification was observed under S stands, compared to B and D (Figure 1b). Tree species composition may affect mineral topsoil acidity and soil exchangeable base cation stocks in various ways. With broadleaf species, e.g., European beech, having markedly larger foliage nutrient and lower foliage lignin contents than conifer species, spruce and Douglas fir (Augusto et al. 2002), they accelerate litter decomposition and bioturbation and promote thin organic layers with comparable high base saturation and less acidity. The introduction of acidic and nutrient-poor litterfall in conifer stands contributes to high concentrations of exchangeable Al in top mineral soil (0-5 cm) (Table S2) under D and S stands, as also reported by (Hansson et al., 2011). This effect was especially clear at the NR sites. The NR sites are characterized by low clay content (Table 1) and high microbial activity compared to SR sites (Lu et al., 2021) and this might accelerate litter decomposition. However, the differences at the SR sites were not consistent with previews studies (Cremer et al., 2017, Augusto et al., 2002). Our results showed that under clay soil conditions, B and D showed more acidic condition than S stands below 5 cm soil depth. Although all plots received the same amount of liming (see Table S1), it is interesting, that B and D stands developed lower pH values (−0.28 units) in the deeper mineral soil than S stands. The acidification in the mineral soil at a depth of 5-30 cm occurs in the main rooting zone typical for B and D (Bolte and Villanueva, 2006; Lwila et al., 2021). These authors showed that B is expected to have about four times higher fine roots biomass than that observed of S stands (Bolte and Villanueva, 2005, Lwila et al., 2021). We did not directly determine the fate of nutrient uptake from deeper mineral soil to biomass formation, however the base pump effect may serve as one relevant explanation for the observed patterns in soil pH. According to the concept of the base pump (Berger et al., 2006), in the beech stands the K^+^, Mg^2+^ and especially Ca^2+^ were taken up via the tree fine root system in exchange of H^+^ (Binkley, 2019; Binkley et al., 2004) and were sequestrated in the tree biomass. This process contributes to mineral soil acidification, as free acid is produced and the buffering capacity (ANC) of the subsoil is depleted (Berger et al., 2006). This bio-acidification (Westman and Jauhiainen, 1998) is known as an important process for decreasing pH values in the mineral soil (Hallbacken and Veranderungen, 1992), and has being reported by several authors (Achilles et al., 2021; Berger et al., 2006; Wittich, 1948). Indeed, decreases in base cations stocks (Ca and Mg) at the mineral soil (5-10) were observed under B forests compared to S stands at the SR sites (Figure 2 and Table S2). This pattern is in line with observations of forest soil acidification in Germany of (Wellbrock and Bolte, 2019) and also reported for beech trees covering a broad geologic gradient site conditions in Southern Germany (Pretzsch et al., 2014).

### Possible effects of tree species involved and their mixture on topsoil stocks of nutrient element cations

A comparative consideration of elemental stocks of Ca, Mg and K listed in Table 4 and for the organic layer down to a soil depth of 30 cm is intended in the given context only to indicate possible effects of the tree species involved or their mixtures on the topsoil in the form of obvious patterns. In this context, the determined absolute values of the stocks for all study sites of both regions and in accordance with the mean values from the comprehensive soil condition survey BZE in the entire region (Lower Saxony) are at the respective lower end of the values determined in the BZE (Evers et al., 2019)

In comparison to e.g., a selected BZE recording point (BZE NI00047; Evers et al., 2019) and considering comparable soil type groups (Triassic sandstone material), the same species and age group (> 70 years), identical (mineral soil: NH4Cl percolation) or nearly identical analytical methods (organic layer samples, digested by HNO3 or aqua regia), the results from the BZE and the given dataset for the total stockpile down to 30 cm soil depth are in the same range (Ca: 136 - 157; K: 99 - 120 kg.ha-1), but also clearly deviating (Mg: 34 - 89 kg ha^-1^). The main reason for the often-high scatter in the comparative determination of nutrient stocks, especially in the topsoil of forest sites, are, despite largely conditioned sample processing and analysis steps, the per se high spatial heterogeneity (see also Table 1 and Table 2). This affects in particular the reliable determination of the mean humus layer amounts, as well as the delimitation of individual horizons, and this, especially between the lowest layer of the organic layer (Oh) and the first mineral soil horizon (Ah).

However, in the present study, these adversities were considered to the greatest possible extent, especially when taking samples in the forest to estimate the topsoil stocks. Thus, all samples were collected and processed exclusively by the same and experienced person and with exactly the same procedures. The same applies to the analytical procedure. Thus, within this study, a comparative observation of the values obtained appears to be permissible.

For the first identification of possible effects by the tree species or their mixtures in the topsoil, the ratio of elemental stocks of Ca, Mg and K exchangeable bound (NH_4_Cl; see method section) in the upper mineral soil (0-30 cm) to the total stocks determined down to a soil depth of 30 cm, i.e., including the stocks of the organic layer determined by means of acid digestion (HNO_3_), was therefore used. The assumption is that such a consideration allows possible conclusions with regard to the material turnover from the litter, respectively the accumulation of nutrients in the organic layer - which is to be evaluated as rather unstable - up to the exchangeable binding in the mineral topsoil - which is regarded as stable. In this context, an increased rooting of the mineral topsoil, as a result of a relatively increased nutrient supply here, seems fundamentally desirable for reasons of ecological stability and a sustainable nutrient supply.

First possible indications in this direction arise with regard to the Ca stocks. Here, there is a clear tendency that under B and D but also their mixture (BD) the relatively highest, and under S, or their mixture with B, the respective lowest percentage stocks in the mineral soil were determined. It is concluded that there is an increased decomposition and release of Ca inputs via litter under B, and under D and their mixture (DB), compared to S. At the same time, neighbouring studies (Lwila et al., 2021) show that rooting under B and D in the mineral topsoil is increased, compared to S.

Also for Mg, the relatively lowest stocks were found in the mineral soil under S or their mixture with B, and this was particularly pronounced in the northern region (NR). Thereby, all forests in the SR sites show significantly higher total Mg stocks. Spruce, in both regions, showed highest total Mg stocks as well as B and its mixture almost exclusively the lowest. The D stands indicated medium total Mg stocks. Again, relatively poor litter decomposition and release of Mg under S stands is inferred, and relatively high release and uptake of Mg under B and its mixture with D might occur.

It must be considered, however, that the tree species involved are known to be systematically characterized by different interception deposition, especially for the elements considered here (conifers > deciduous trees; S ≥ D > B; De Schrijver et al., 2007) and that this influence cannot be directly accounted in our study. However, this may explain the highest overall accumulation for Mg, combined with the relatively highest proportions in the organic layer, in both regions under S stand.

The picture for K is less clear in our study. Here, due to the initial substrate (loess over Triassic sandstone), systematically higher total stocks are found in the Solling region (SR) and also here, as for Mg, the relatively highest accumulations were determined in the organic layer under S. The same applies to the NR site. Again, and as similar to Mg, it is concluded that there is a reduced litter turnover with subsequent K release under S stands. However, otherwise in the northern region the stocks and their distributions are similar at a low level between the variants, which does not allow further conclusions. What is noticeable, on the other hand, is that in both regions relatively high proportions of K stocks are found in the mineral soil under B and the highest proportions under the mixture of B and D, respectively. A possible cause is again a more intense litter turnover under B, compared to S, with a subsequent binding in the upper mineral soil. This process is possibly further promoted by the admixture of D.

Moreover, higher CEC in the mineral topsoil under D stand at the NR sites can be attributed to higher contents and stocks of soil organic carbon (SOC) (Foltran, *in preparation*) due to the importance of soil organic matter (SOM) in cation. Cremer and Prietzel, (2017) ascribed the higher CEC of SOM to conifer stands in comparison to broadleaves, supporting our findings of higher CEC in D stands topsoils compared to pure B. The SOC contents differ among species at the present sites (Foltran, *in preparation*) and are generally higher under conifers than B. Top-mineral soil (0-30 cm) OC contents were positively correlated with topsoil CEC (*p* =0.83) and negatively correlated with topsoil pH (*p*=-0.4) (Figure S1). Tree species impacts on CEC are likely to be indirect effects, triggered by the effect of tree species on topsoil OC parameters.

### Beech and spruce depletes soil Ca and Mg, but Douglas fir depletes soil K stocks

The Douglas fir and spruce stands effects on mineral soil acidity (described above), also showed effects on total soil exchangeable Ca and Mg stocks (Figure 2). In both regions the highest total soil exchangeable Ca and Mg stocks were observed under the D stand, mirrored the high Ca and Mg stocks observed on the organic layer (Figure 2 and 3).

Highest total soil exchangeable Ca and Mg storage in Douglas fir stands may reflect the patterns of nutrient uptake and accumulation in tree biomass by tree species. Nutrient contents of wood can provide a reliable estimation of net nutrient uptake by tree species and nutrient export due to wood harvesting in managed forests. Despite of its highest biomass production, Douglas fir trees are characterized by lowest base cation contents, originating mainly from particularly low contents of Ca, Mg and K in wood and bark compared to spruce and especially beech (Augusto et al. 2000; Pretzsch et al. 2014). Overall, nutrient immobilization in aboveground biomass of fast growing species, i.e., Douglas fir stands with highest tree biomass (deduced from highest values of basal area and tree height; Table 2) compared to beech or Norway spruce, thus preserving soil nutrient stocks more than in pure beech and spruce stands, where nutrient removal of Ca and Mg are showed to be the highest (Pretzsch et al. 2014).

Confirming earlier results of Mareschal et al. (2010) and Cremmer et al., (2017), mineral topsoil K concentrations and stocks (Table S1; Figure 2), in contrast to Ca and Mg, were higher in beech compared to Douglas fir stands. Under beech, high mineral soil exchangeable K stocks (0-10 cm) compensate for smallest organic layer (L-layer) K stocks (Figure 3). Canopy exchange of base cations, as the major source for K input via throughfall deposition, is higher in deciduous stands compared to conifer forests (De Schrijver et al. 2007), compensating for higher dry deposition in the latter. Coincidentally higher K contents in beech leaves compared to Douglas fir and spruce needles (Pretzsch et al. 2014; Cremeretal. 2016) might furthermore balance K inputs between beech and the conifers.

### Site dependent differences in topsoil acidity between Douglas fir and spruce

At the NR sites, high acidity and low BS under S stands than D can be caused by higher Mg and K contents in D needles compared to S (Pretzsch et al. 2014; Cremer et al. 2016) and likely promote nutrient return into the soil under D compared to S stands. Moreover, differing rooting patterns among D and S might have contributed to the differentiation between the conifers. The restriction of S roots to the forest floor and the uppermost mineral soil (Lwila et al., 2021), while D also penetrates deeper soil layers (Spielvogel et al. 2014), reinforces soil acidification in S topsoil as observed in our study at sandy soils (NR sites). However, we can only hypothesize possible causes for topsoil differences in pH among D and S at the SR sites. There, different than expected, S stands presented less acidic conditions than D stands at 10-30 cm mineral soil depth. It is important to point that pH differences between Douglas fir and spruce are null at 0-5 cm. However, below 5 cm the differences are significant (p>0.05), ranging from −0.31 units (5-10 cm) to −0.25 units (10-30 cm). Possible explanations can be either previous land use, as former spruce stands are likely to be replaced by other conifer, i.e., Douglas fir, or bio-acidification as described above. Our results showed that D likely deplete K, but no changes on Ca and Mg storages were observed. In fact, under D stands high stocks of base cations were observed suggesting that D might improve soil nutrition, independent of site conditions. Large quantities of nutrients are stored in the organic layer (Table S2) and leading to an improvement of nutrient cycle, especially important under sandy soil conditions as NR sites.

### Tree species mixtures effects on mineral topsoil

According to our results, admixture of beech to Douglas fir, and especially to spruce, counteracts soil acidification by increasing mineral topsoil pH and BS under loamy soil (17.7% – DB; 24.2% – SB, SR sites) and sandy soils (25.0% – DB; 19.0% – SB, NR sites). The same effects were observed by Berger et al. (2009), where soils in mixed beech-spruce stands tended to have higher BS than pure spruce.

Our results indicate that at the NR sites (sandy soils) beech–Douglas fir mixtures can be superior to beech–spruce mixtures with respect to BS, CEC and exchangeable K and tended to be less acidic. Under this condition, litter decomposition and bioturbation are favoured, thus reducing the amount of nutrient immobilisation (e.g. Ca and K) in the organic layer (Figure 2). Similarly to pure spruce at SR sites, and in contrast to DB mixtures, the SB forest tended to exploit mineral soil exchangeable Ca, Mg and K stocks.

Overall, our data provides basic knowledge on DB mixtures and their impact on European forest soils compared to adjacent pure stands and mixed stands of beech with Norway spruce. Generally, species mixtures effects on organic layer and mineral topsoil acidity and base cations were additive effects, meaning that DB and SB mixtures were within the range corresponding to the pure stands. There were only few and mostly inconsistent (across sites and species mixtures) deviations from this pattern, suggesting that there were hardly any non-additive effects of species mixtures. Though mixed stands are often assumed to be intermediate in comparison with the pure stands (additive effects) of the respective species concerning their soil chemical properties (Rothe and Binkley 2001), this strongly depends on site condition i.e., loamy vs sandy soils, and the specific tree species involved (Augusto et al. 2002). Confounding effects of differing environmental and stand parameters may modify trends and magnitudes of tree species mixtures effects (Richards et al. 2010). Thus, underlying mechanisms still need to be explored in more detail.

## Conclusions

Our assumptions that i.) admixing Douglas fir to beech forests increases nutrient availability, and that ii.) the nutrient pool of Douglas fir and beech monocultures are comparable but differ from Norway spruce, were confirmed by our data. Soil exchangeable Ca and Mg stocks in Douglas-fir and European beech forests were significantly higher than in Norway spruce stands. Moreover, we hypothesized that iii.) under reduced nutrient availability, species-identity effects will be stronger expressed, compared to more rich soils. Indeed, mixed Douglas-fir-beech showed expressive differences at Northern sites.

Overall, our study suggest that the enrichment of beech stands by Douglas fir does not cause unexpected and detrimental changes of soil acidity and does not strongly affect soil exchangeable base cation when compared to pure beech. Instead, admixtures of Douglas-fir seem to lead to smaller changes in pH, CEC and BS than those of Norway spruce. Therefore, forest management may consider mixtures of European beech and Douglas fir as a reasonable management option without apprehending negative effects on soil chemistry.

## Author contributions

All authors contributed to the study conception and design. Material preparation, data collection and analysis were performed by Estela Foltran. The first draft of the manuscript was written by Estela Foltran and all authors commented on previous versions of the manuscript. All authors read and approved the final manuscript.

## Acknowledgements

The study was conducted as part of the Research Training Group 2300 funded by the German research funding organization (Deutsche Forschungsgemeinschaft – DFG, contract number 316045089, SP3). We gratefully acknowledge the administrative support by Serena Müller and the indispensable help of Julian Meyer and Dirk Böttger during soil sampling. Furthermore, we thank Sylvia Bondzio, Karin Schmidt for their valuable advice during laboratory work. We also thank Jingzhong Lu for the fruitful discussions about the statistical analysis and Dr. Ulrike Talkner (North-West German Forest Research Centre) for providing the liming dataset from Lower Laxony.

---

**Table S 1.** Liming data

| Site | Plot | Species | Year | CaCO<sub>3</sub> (Mg ha<sup>-1</sup>) | MgCO<sub>3</sub> (Mg ha<sup>-1</sup>) | P<sub>2</sub>O<sub>5</sub> (Mg ha<sup>-1</sup>) |
| --- | --- | --- | --- | --- | --- | --- |
| Solling (SL) | Dassel | D | 2010 | 1.65 | 1.05 | 0 |
|  |  | DB | 2010 | 1.65 | 1.05 | 0 |
|  |  | S | 1991 | 2.25 | 0.45 | 0 |
|  | Winnefeld | D | 2008 | 1.57 | 1.25 | 0 |
|  |  | DB | 2008 | 1.5 | 1.19 | 0 |
|  |  | B | 2008 | 1.57 | 1.25 | 0 |
|  |  | S | 2008 | 1.57 | 1.25 | 0 |
|  | Nienover | D | 2008 | 1.57 | 1.25 | 0 |
|  |  | DB | 1995 | 1.5 | 1.2 | 0 |
|  |  | B | 1993 | 1.65 | 1.05 | 0 |
|  |  | B | 2008 | 1.57 | 1.25 | 0 |
|  |  | SB | 1993 | 1.65 | 1.05 | 0 |
|  |  | SB | 2008 | 1.57 | 1.25 | 0 |
|  |  | S | 1993 | 1.65 | 1.05 | 0 |
|  |  | S | 2008 | 0.62 | 0.49 | 0 |
| Northern (NS) | Nienburg | D | 2001 | 1.8 | 0.9 | 0 |
|  |  | DB | 1990 | 1.5 | 1.2 | 0 |
|  |  | DB | 2001 | 1.8 | 0.9 | 0 |
|  |  | S | 1990 | 1.48 | 1.19 | 0 |
|  |  | S | 2001 | 1.8 | 0.9 | 0 |
|  | Unterlüß | DB | 1990 | 1.4 | 0.12 | 0.47 |
|  |  | B | 1990 | 1.4 | 0.12 | 0.47 |
|  |  | SB | 1990 | 2.17 | 0.37 | 0.09 |
|  | Göhrde II | No liming recorded in the last 30 years |  |  |  |  |
|  | Göhrde I | No liming recorded in the last 30 years |  |  |  |  |

**Table S 2.**
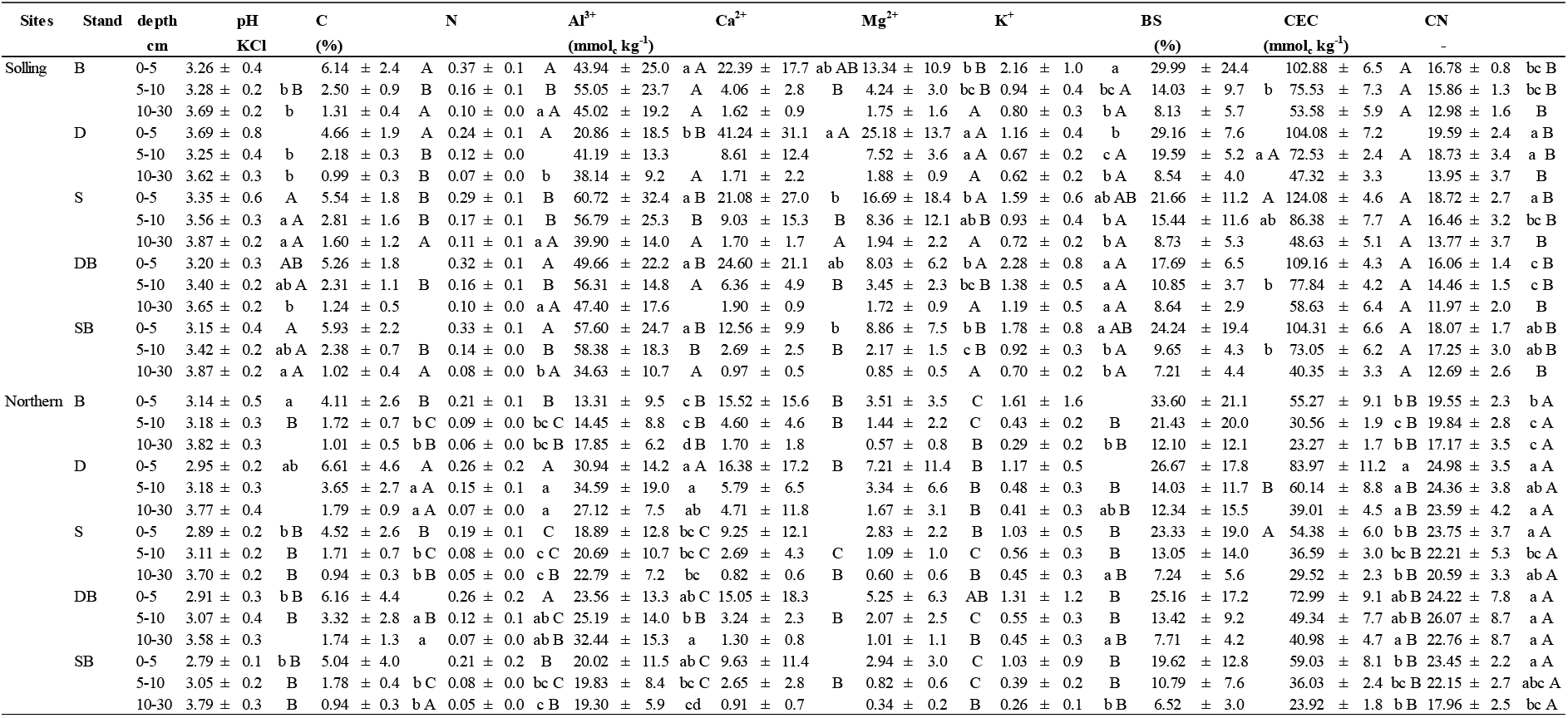
Soil pH (KCl), C and N concentration, exchangeable Al, Ca, Mg, K concentration (mmolc kg^-1^), base saturation (BS; %), cations exchangeable capacity (CEC; mmol_c_ kg^-1^) and C:N ratio in various depths of the mineral soil of pure stands of Douglas fir (D), Norway Spruce (S), European beech (B) and mixed stands Douglas fir + Beech (DB) and Norway spruce + Beech (SB). Average values and standard deviation are presented by stand type and region, Solling (*SR, n*=12), and Northern (*NR, n*=16). Significant (p < 0.05; Kruskal-Wallis-H-test, followed by pairwise Mann-Whitney-U-test) differences between tree species for a given soil depth are marked by different letters: lowercase letters for individual soil depths inner sites and capitals between sites for the same species.

**Table S 3.**
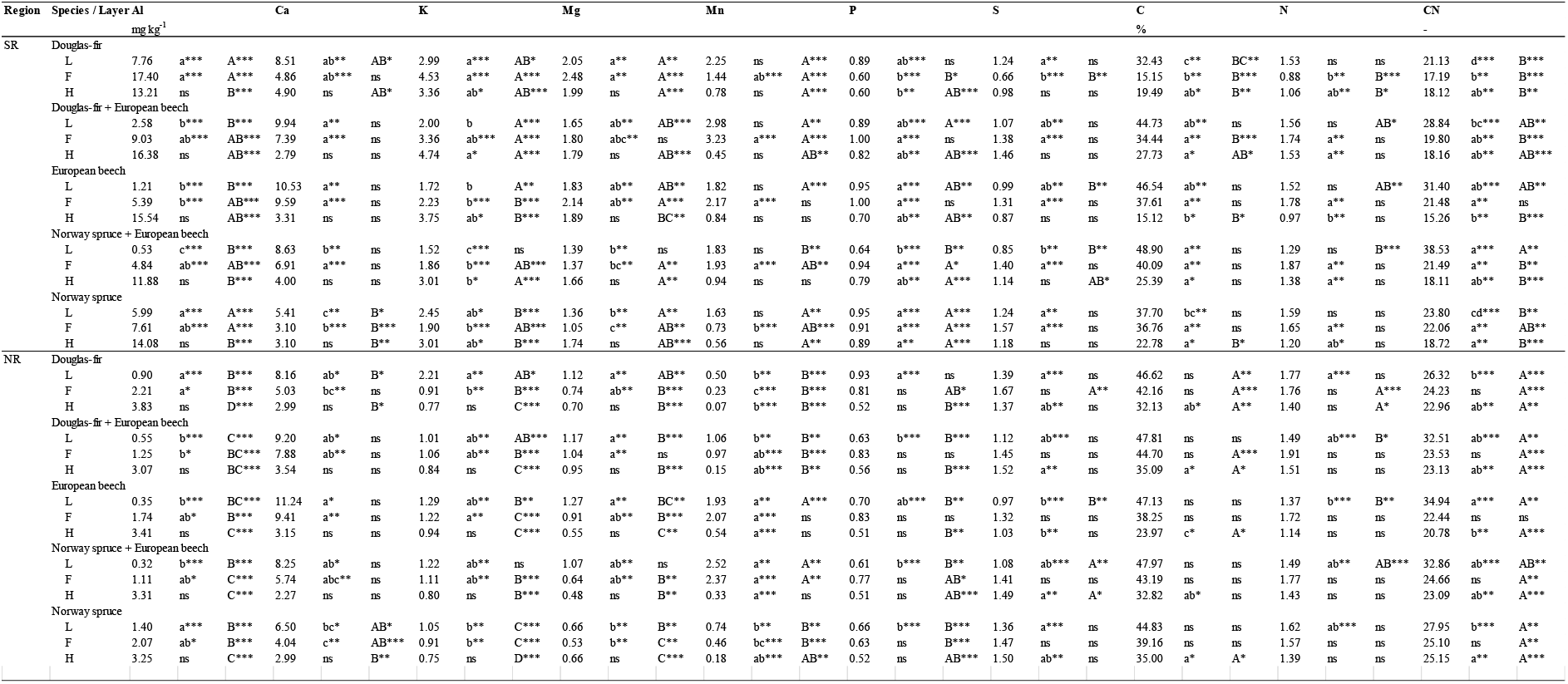
Nutrient concentration in various layer of organic layer of pure stands of Douglas fir (D), Norway Spruce (S), European beech (B) and mixed stands Douglas fir + Beech (DB) and Norway spruce + Beech (SB). Average values and standard deviation are presented by stand type and region, Solling (*SR, n*=12), and Northern (*NR, n*=16). Significant (p < 0.05; Kruskal-Wallis-H-test, followed by pairwise Mann-Whitney-U-test) differences between tree species for a given soil depth are marked by different letters: lowercase letters for individual soil depths inner sites and capitals between sites for the same species.

**Figure S 1.**
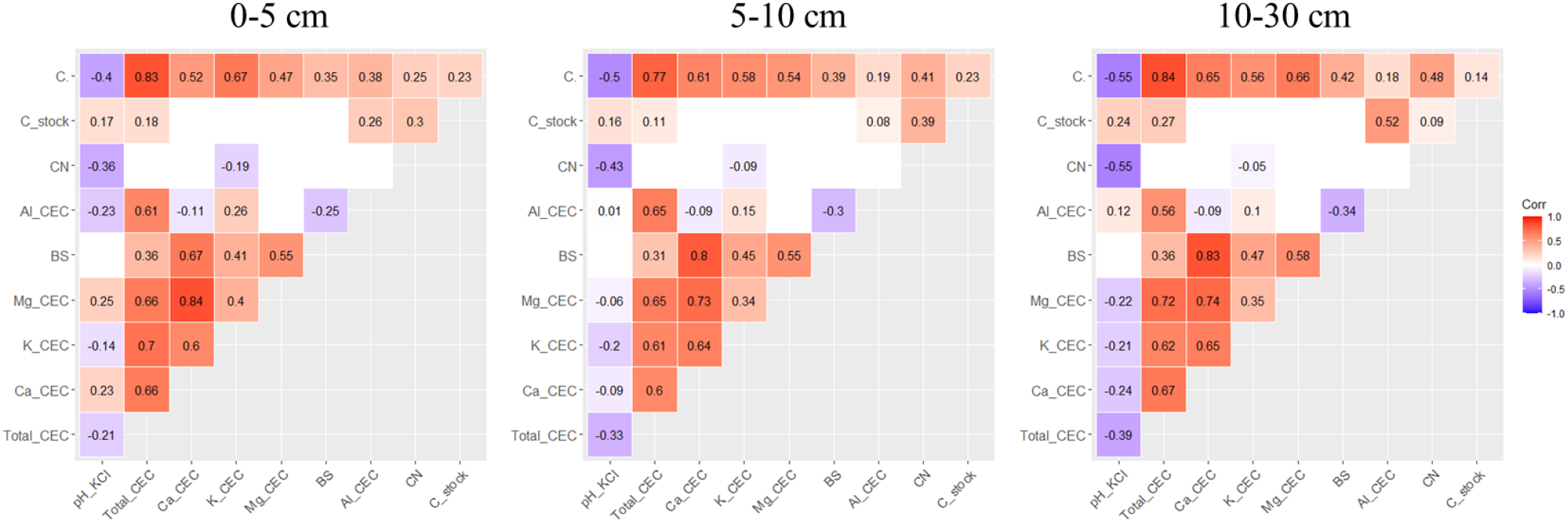
Spearman correlation among mineral soil variables, C content, C stocks, C:N ratio, exchangeable Al^3+^,Mg^2+^, K^+^, Ca^2+^ and total CEC for each mineral soil depth (cm). Significant correlations are marked in bold.

## Notes

### Competing Interest Statement

The authors have declared no competing interest.

